# MAGGIE: leveraging genetic variation to identify DNA sequence motifs mediating transcription factor binding and function

**DOI:** 10.1101/2020.01.30.925917

**Authors:** Zeyang Shen, Marten A Hoeksema, Zhengyu Ouyang, Christopher Benner, Christopher K Glass

## Abstract

Genetic variation in regulatory elements can alter transcription factor (TF) binding by mutating a TF binding motif, which in turn may affect the activity of the regulatory elements. However, it is unclear which TFs are prone to be affected by a given variant. Current motif analysis tools either prioritize TFs based on motif enrichment without linking to a function or are limited in their applications due to the assumption of linearity between motifs and their functional effects. Here, we present MAGGIE, a novel method for identifying motifs mediating TF binding and function. By leveraging measurements from diverse genotypes, MAGGIE uses a statistical approach to link mutation of a motif to changes of an epigenomic feature without assuming a linear relationship. We benchmark MAGGIE across various applications using both simulated and biological datasets and demonstrate its improvement in sensitivity and specificity compared to the state-of-the-art motif analysis approaches. We use MAGGIE to reveal insights into the divergent functions of distinct NF-κB factors in the pro-inflammatory macrophages, showing its promise in discovering novel functions of TFs. The Python package for MAGGIE is freely available at https://github.com/zeyang-shen/maggie.

## 1. Introduction

Genome-wide association studies (GWAS) have identified thousands of genetic variants associated with an increase in disease risk (MacArthur et al., 2017; Visscher et al., 2017). The majority of these variants fall within regulatory elements such as promoters and enhancers, implicating an effect on transcriptional regulation (Khurana et al., 2016; Farh et al., 2015; GTEx Consortium, 2015). Transcription factors (TFs) play an essential role in mediating the activity of regulatory elements. Many TFs possess DNA-binding domains that recognize specific DNA sequences, called TF binding motifs. Genetic variation that alters TF binding motifs has been established as an important mechanism for regulatory variants to affect transcriptional regulation (Grossman et al., 2017; Deplancke et al., 2016; Heinz et al., 2013). However, it is not always straightforward which TF is affected by a given variant. First of all, a genetic variant is able to alter multiple motifs. Binding motifs for hundreds of TFs are currently available in the public databases (Fornes et al., 2020; Weirauch et al., 2014; Matys et al., 2006). Many motifs correspond to similar or overlapping DNA sequences, which can be altered by the same variant simultaneously. The second complication is due to the great dependency of TF binding on conditions. Multiple TF binding motifs are usually packed at regulatory elements across 100-200 base pairs (Lambert et al., 2018; Tsompana and Buck, 2014) but can become functional under different conditions depending on cell type, stimulus, time point, etc. (Eeckhoute et al., 2009; Spitz and Furlong, 2012) Knowing the function of motifs for a given condition can help prioritize TFs prone to be affected by genetic variation and ultimately have an impact on transcriptional regulation.

Numerous motif analysis tools have been published in the past decade in order to prioritize important TFs for future validation (Jayaram et al., 2016; Boeva, 2016). One major category of tools identify enriched motifs that appear more frequently at given regions of interest than random genomic regions (Heinz et al., 2010; Machanick and Bailey, 2011; Siebert and Söding, 2016). Due to the development of high-throughput sequencing assays, these approaches can now be ap-plied to various types of epigenomic features, such as chromatin accessibility measured by the assay for transposase-accessible chromatin using sequencing (ATAC-seq) or DNase I hypersensitive sites sequencing (DNase-seq), and TF binding and histone modification measured by chromatin immunoprecipitation sequencing (ChIP-seq), etc. (Reuter et al., 2015). However, motifs prioritized by enrichment algorithms may include many false positives, which have no impact on the epigenomic feature of interest if being mutated, because, as previously discussed, enriched motifs are not necessarily bound by TFs or functional for an epigenomic feature under a given condition.

Another category of motif analysis tools prioritize TFs by leveraging measurements and genetic variation of human individuals or animal strains, such as MMARGE (Link et al., 2018b) and TBA (Fonseca et al., 2019). Both methods depend on an assumption of linearity between motifs and epigenomic features. This assumption worked for TF binding but likely does not hold for many other epigenomic features like histone modification or stimulus response of regulatory elements. These features result from the interactions between many TFs and may not possess a simply linear relationship with TF binding motifs.

Here, we developed a novel approach, MAGGIE (Motif Alteration Genome-wide to Globally Investigate Elements), to identify DNA motifs mediating TF binding and function. Considering the increasing measurements of genotypes and various epigenomic features for different individuals and animal strains, we are able to identify genomic regions differentially associated with some epigenomic feature of interest across different genotypes, labeling them as positive or negative for sequences with or without the feature, respectively (Fig. 1A). We propose to associate these differential regions with changes of TF binding motifs caused by genetic variation to gain insights into the functions of motifs. Unlike conventional motif enrichment methods, MAGGIE is independent of the background frequency of motifs and gains power in picking up functional motifs by leveraging motif mutations at the same regions between individuals or strains. MAGGIE differs from MMARGE and TBA by eliminating the assumption of linearity between motifs and testing features. This design of the framework is flexible in accommodating any type of epigenomic feature, including but not limited to the ones to be discussed in this paper, such as TF binding, open chromatin, histone modification, and stimulus response of regulatory elements.

**Fig. 1.**
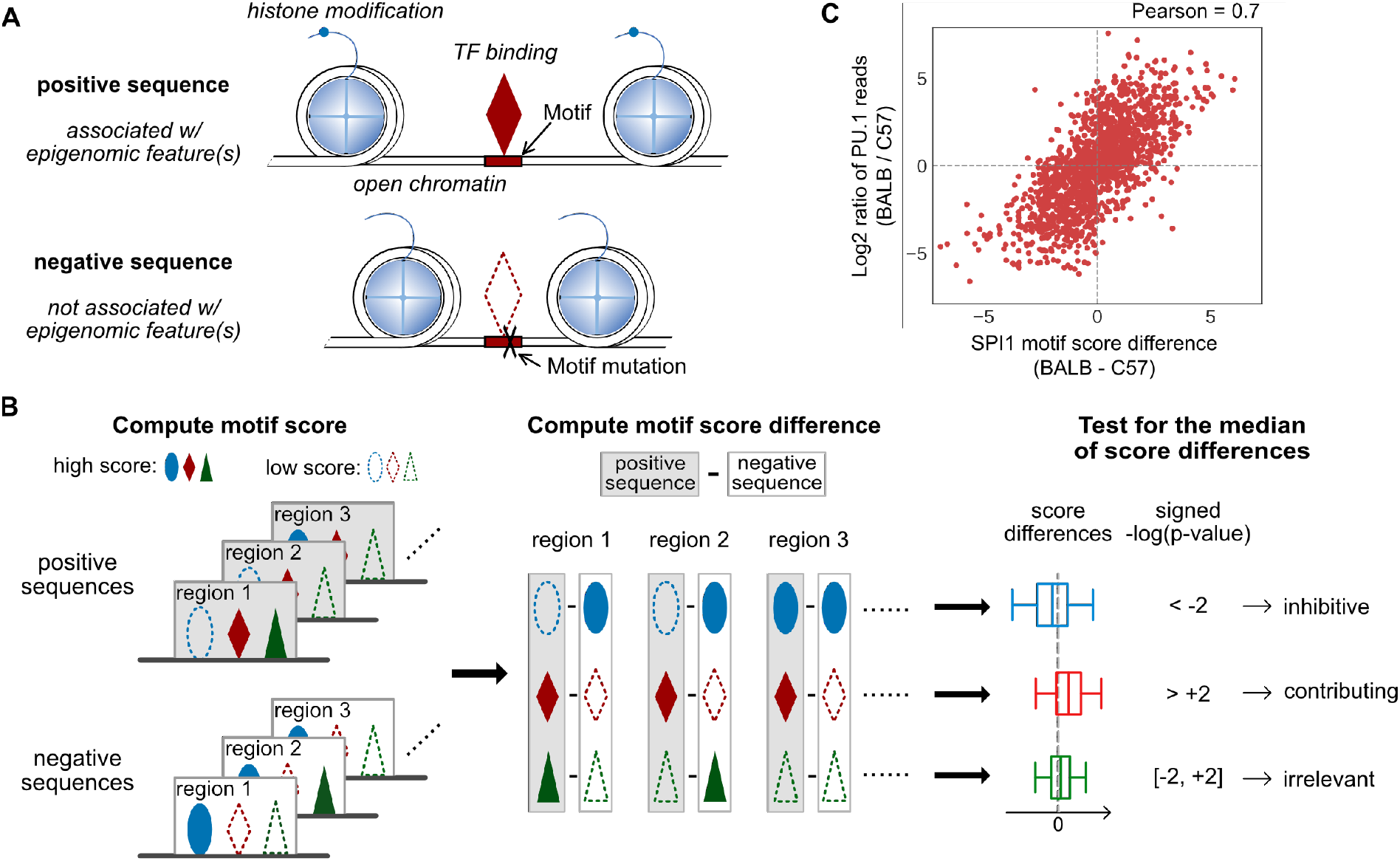
The schematics of MAGGIE. (**A**) Schematic depicting how the epigenomic features of regulatory elements are related to the inputs of MAGGIE. Positive sequences are defined to be associated with one or more epigenomic features of interest, such as TF binding, open chromatin, histone modification, etc. Each positive sequence is paired with a counterpart negative sequence, which has a loss of the chosen epigenomic feature(s) due to mutations on certain TF binding motifs. (**B**) Flowchart of MAGGIE. Positive and negative sequences are input to compute motif scores as an estimated likelihood of being bound by certain TF. A representative motif score is obtained for each sequence by taking the maximum, displayed by different shapes (ellipse, diamond, and triangle) for different TFs. High motif scores are shown as solid shapes and low scores as dashed shapes. Next, differences of representative motif scores are computed for every TF by subtracting scores of negative from positive sequences. Finally, the score differences for each TF are aggregated, and the median value is tested by Wilcoxon signed-rank test to evaluate whether there is a bias in the changing direction from positive to negative sequences. The examples demonstrate a significant bias of increase (ellipse) or decrease (diamond) or an insignificant bias (triangle), which implicate the inhibitive, contributing, or irrelevant roles of TFs, respectively. (**C**) Correlation between differences of representative motif scores for SPI1 motif and log2 fold changes of PU.1 binding activity between BALB and C57 mice. The score differences and the log fold changes were calculated by subtracting C57 from BALB.

We evaluated the performance of MAGGIE in both simulated datasets and biological datasets and compared our results to two other approaches, HOMER (Heinz et al., 2010) and MMARGE (Link et al., 2018b), which are representative for the two major categories of motif analysis tools discussed before. The results demonstrated the superior sensitivity and specificity of MAGGIE in all of the experiments. By applying MAGGIE to the regulatory elements of macrophages in response to pro-inflammatory stimulus, we captured divergent functions of distinct NF-κB factors despite the similarity of their motifs. These results were further validated by the NF-κB binding sites measured by ChIP-seq experiments, showing the promise of MAGGIE in identifying highly specific motifs and discovering novel functions of TFs.

## 2. Materials and methods

### 2.1 Overview of MAGGIE

The overall framework of MAGGIE is illustrated in Figure 1B. MAGGIE takes pairs of sequences as inputs. Positive sequences are identified to be associated with some epigenomic feature of interest, while negative sequences are from different alleles or the same regions of a different genome where the chosen epigenomic feature is not found. Depending on the genetic difference of genomes, every pair of input sequences can have a variable number of genetic variants like single nucleotide polymorphisms (SNPs) and short insertions and deletions.

The basic assumption for MAGGIE is that the allele specificity of epigenomic features is derived from the genetic variation between positive and negative sequences that mutate certain TF binding motifs. This assumption is supported by the findings that motif mutations due to local genetic variation is the major explanation for the gain or loss of TF binding sites (Link *et al.*, 2018a; Kundaje *et al.*, 2015). Considering the importance of transcription factors for other epigenomic features like promoter and enhancer function (Reiter *et al.*, 2017; Spitz and Furlong, 2012), we hypothesized that our framework could help identify motifs mediating both TF binding and other epigenomic features affected by TF binding.

The computation of MAGGIE is centered on the motif score based on position weight matrix (PWM), which is the widely used metric to approximate the likelihoods of being bound by certain TF (Stormo, 2000). Given pairs of positive and negative sequences associated with a chosen allele-specific epigenomic feature, MAGGIE computes motif scores for hundreds of TFs whose PWMs are currently available in the JASPAR database (Fornes *et al.*, 2020). For each TF, a representative motif score is calculated for every sequence by taking the maximum score across the sequence. MAGGIE then computes differences of representative motif scores by subtracting scores of negative from positive sequences to obtain the changes of binding likelihood. Score differences should have a bias towards positive values (i.e., higher motif scores in positive sequences) if the corresponding TF is contributing to the chosen epigenomic feature. On the contrary, if the TF is potentially inhibitive for the chosen feature, the aggregated differences will tend to have negative values (i.e., lower motif scores in positive sequences). Irrelevant TFs will have their motifs randomly mutated by genetic variation, so the score differences should be overall balanced around zero. A nonparametric Wilcoxon signed-rank test is used to statistically test the significance of the association between motif mutations and the chosen epigenomic feature by asking whether the median of all the non-zero motif score differences is close to zero. A signed p-value showing the sign of the median value together with the p-value from the statistical test implicates the function of TF to be either contributing (positive) or inhibitive (negative) if being called significant.

### 2.2 Computation of motif score and motif score difference

Motif score is a reliable metric to measure the likelihood of TF binding and can well reflect the binding activity of the corresponding TF (Boeva, 2016; Ji *et al.*, 2018). A position weight matrix (PWM) stores the log likelihoods for the four possible nucleotides (A, C, G, and T) to be bound by a TF at each position (Stormo, 2000):

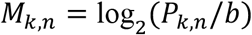

where *P*_*k,n*_ is the probability of seeing nucleotide *n* at the *k*^th^ position of the motif, obtained from the measurements of ChIP-seq, protein binding microarrays, HT-SELEX, etc. (Lambert *et al.*, 2018), and *b* is the background probability, usually set as 0.25. Given a DNA sequence, we can compute motif scores for any of those TFs by adding up the log likelihoods of seeing certain nucleotides at every position:

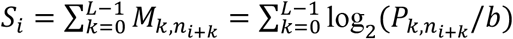

where *S*_*i*_ is the motif score for a segment of the given sequence from position *i* to position *i*+*L*−1, supposing *L* is the length of the motif and *i* starts at 1, and *n*_*i*+*k*_ is the nucleotide at position *i*+*k*. For a sequence longer than the motif (i.e., the biggest possible *i* greater than *L*), instead of dealing with a list of motif scores, we obtained the maximum motif score to represent the binding likelihood of the complete sequence:

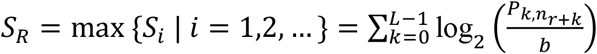

where *r* is the starting position of the maximum motif score. For any pair of positive and negative sequences, we ended up with two representative motif scores:

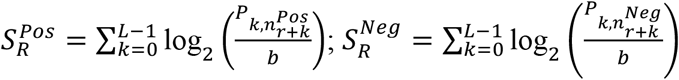

Then, the log fold change of binding likelihood within the sequence pair can be computed by subtracting the representative motif score of the negative sequence from that of the positive sequence:

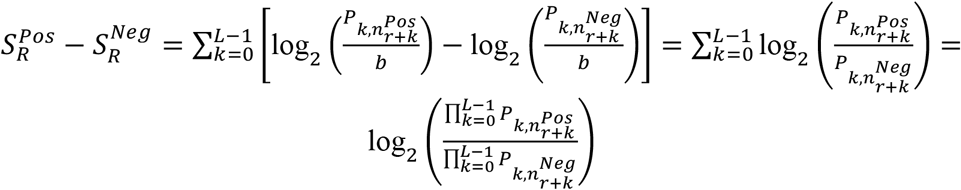

The difference of representative motif score turns out to be the log fold change of the binding likelihood between positive and negative sequences. Any representative motif score less than zero is replaced by zero before the computation of score difference, reducing impacts from poorly matched motifs. The difference of representative motif score is a good indicator of the binding activity change of the corresponding TF (Martin *et al.*, 2019; Spivakov *et al.*, 2012). For example, by comparing PU.1 binding in macrophages of C57BL/6J (C57) and BALB/cJ (BALB) mice (Link *et al.*, 2018a), we observed a strongly positive correlation between the score difference of PU.1 (or SPI1) motif and the log fold change of PU.1 binding activity quantified by ChIP-seq reads among 1635 PU.1 binding sites that have SPI1 motif mutations between the two mouse strains (Fig. 1C). In most cases, positive score differences, equivalent to a higher binding likelihood, result in stronger binding. A larger scale of score difference often corresponds to a larger change in TF binding, and this relationship is independent of the actual motif score (Supplementary Fig. S1). These intrinsic characteristics of motif score difference support the hypotheses that i) motif score difference can indicate change of binding of the corresponding TF, and ii) aggregated motif score differences can reflect whether the presence of the chosen epigenomic feature is associated with the gain or loss of TF binding.

### 2.3 Applications and data preparation

#### 2.3.1 Simulated data

To characterize the performance of MAGGIE and systematically compare to other methods, we generated simulated positive and negative sequences. Positive sequences are generated by first randomly selecting the four possible nucleotides to form sequences of 200-base pair (bp). Then we create TF binding motifs by sampling nucleotides based on their probabilities derived from PWMs and insert these motifs at non-overlapping positions. To obtain counterpart negative sequences, SNPs are simulated inside hypothetic “contributing” motifs by switching the existing nucleotides to other nucleotides.

During the generation of simulated data, we can insert “irrelevant” motifs, which experience either no mutation or random mutation, to evaluate the specificity of MAGGIE. The sensitivity of MAGGIE can be tested by changing the number of simulated sequences (i.e., sample size) or the fraction of sequences having motif mutations (i.e., signal-to-noise ratio).

#### 2.3.2 Transcription factor binding sites

We first tested MAGGIE to identify TF binding motifs for corresponding TF binding. Allele-specific binding sites of 12 TFs were obtained from two cell types, GM12878 and HeLa-S3 (Shi et al., 2016). We extracted 100-bp sequences around the SNPs associated with allele-specific binding sites and labeled the sequences with the binding alleles as positive sequences and those with the non-binding alleles as negative.

MAGGIE was then used to identify collaborative TFs. We downloaded the ChIP-seq data of PU.1 and C/EBPβ for C57 and BALB mice from the GEO database with accession number GSE109965 (Link et al., 2018a), and the ChIP-seq data of ATF3 for the same mouse strains with the accession number GSE46494 (Fonseca et al., 2019). The data for C57 were mapped to the mm10 genome using Bowtie2 v2.3.5.1 (Langmead and Salzberg, 2012), while the data for BALB were first mapped to the BALB genome and then shifted to the mm10 genome using the MMARGE v1.0 “shift” function (Link et al., 2018b). The reproducible TF binding sites were identified by using HOMER v4.9.1 to call unfiltered 200-bp peaks (Heinz et al., 2010) and running IDR v2.0.3 on replicates with the default parameters (Qunhua Li and Bickel, 2011). The TF binding sites found only in one of the strains were defined to be strain-specific, yielding 13099 PU.1, 8127 C/EBPβ, and 13347 ATF3 strain-specific binding sites between BALB and C57. The sequences of strain-specific binding sites were extracted as pairs from both strains using the MMARGE v1.0 “extract_sequences” function (Link et al., 2018b). Sequences associated with TF binding are labeled as positive regardless of which strain they are originated from, and their counterpart sequences from the other strain are labeled as negative.

#### 2.3.3 Chromatin quantitative trait loci

The application of MAGGIE was further extended to discover motifs mediating chromatin accessibility and histone modification. DNase I sensitivity quantitative trait loci (dsQTLs) were downloaded from the GEO database with accession number GSE31388 (Degner et al., 2012). Histone quantitative trait loci (hQTLs) were acquired for three types of histone modifications, local acetylation of histone H3 lysine 27 (H3K27ac), mono-methylation of histone H3 lysine 4 (H3K4me1), and tri-methylation of histone H3 lysine 4 (H3K4me3) (Grubert et al., 2015). All QTLs were originally analyzed using lymphoblastoid cell lines (LCLs) from ~70 individuals. We filtered the tested hQTLs to those with a p-value less than 1e-6 and a distance to their associated peaks less than 1000 bp. After filtering, we ended up with 5705 dsQTLs, 2452 H3K27ac QTLs, 4012 H3K4me1 QTLs, and 1752 H3K4me3 QTLs. Similar to the preprocessing for the allele-specific binding sites, we extracted 100-bp sequences centering around the variants and labeled the alleles associated with a higher trait level as positive and the other alleles as negative.

#### 2.3.4 Stimulus responses of regulatory elements

We used MAGGIE to analyze the stimulus response of regulatory elements, an epigenomic feature that no existing tools have been reported to work for. We downloaded ATAC-seq and H3K27ac ChIP-seq data from macrophages at both basal state and pro-inflammatory state induced by 1-hour treatment of the TLR4-specific ligand Kdo2 lipid A (KLA) from four diverse strains of mice: C57BL/6J (C57), NOD/ShiLtJ (NOD), PWK/PhJ (PWK), and SPRET/EiJ (SPRET) (Link et al., 2018a). Similar to the preprocessing of ChIP-seq data for TFs, the raw reads were mapped and shifted to the mm10 genome. Based on ATAC-seq data, we obtained 200-bp reproducible open chromatin and filtered for intergenic and intronic regions based on HOMER annotations to obtain potential enhancers (Heinz et al., 2010). Open chromatin regions of the two conditions from the same strain were merged and extended from 200 bp to 1000 bp to quantify their activity by the count of H3K27ac ChIP-seq reads. We filtered for active regulatory elements (greater than 16 reads in at least one condition; Supplementary Fig. S4) and computed the change of activity from basal to KLA-treated condition by the fold change of reads. Regions showing a higher or lower level of H3K27ac larger than 2.5-fold after KLA treatment were labeled as “activated” or “repressed”, respectively (Fig. 4A), and those with less than 40% change were labeled as “neutral”. Based on pairwise comparisons across the four mouse strains, regulatory elements labeled as “activated” or “repressed” only in one of the compared strains were called strain-specific and were pooled for analysis.

### 2.4 Comparative methods

In order to evaluate the performance of MAGGIE, we systematically designed a series of analyses to compare MAGGIE with existing methods. MMARGE (Link et al., 2018b) is a recent approach that fits a linear model between motif score and TF binding activity. MMARGE v1.0 was downloaded from https://github.com/vlink/marge. Since MMARGE does not directly work with binary-labeled datasets (e.g., simulated data, QTLs, allele-specific binding sites), as the replacement, we fit a linear model between motif scores and binary labels using statsmodels package (Seabold and Perktold, 2010) for the simulated data, which should reflect the general performance of using a linear model in these tasks.

Another category of motif analysis tool is based on motif enrichment algorithms, such as HOMER (Heinz et al., 2010), MEME Suite (Machanick and Bailey, 2011), BaMM (Siebert and Söding, 2016), etc. We expect any one of these methods to be representative for the others, so we picked HOMER for all the experiments, which was downloaded from http://homer.ucsd.edu/homer/data/software/homer.v4.9.1.zip and run with the default parameters.

### 2.5 Validation experiment

Bone marrow was isolated from C57 mice and differentiated for 7 days using media containing M-CSF to generate bone marrow-derived macrophages (BMDMs) as described previously (Link et al., 2018a). BMDMs were maintained at basal conditions or treated with KLA for 1 hour. For p65 (Santa Cruz, sc-372X) and p50 (Abcam, ab32360) ChIP-seq experiments, 8 million untreated or KLA-treated BMDMs per assay were double-crosslinked using disuccinimidyl glutarate (DSG) and formaldehyde (FA). ChIP-seq was performed using 2 μg of antibody as described previously (Heinz et al., 2018). ChIP DNA was prepared for sequencing using the NEBNext Ultra II DNA library prep kit (NEB, E7645) and sequencing was performed on the HiSeq4000 (75bp SR, Illumina). The binding sites of p65 and p50 were identified using HOMER “findPeaks -size 200” (Heinz et al., 2010) and then merged to obtain co-binding sites and p65- or p50-only binding sites. The binding activity of p65 and p50 was quantified by the count of ChIP-seq reads.

## 3. Results

### 3.1 MAGGIE shows superior specificity and sensitivity on simulated datasets

To evaluate the performance of MAGGIE, we stochastically simulated one thousand DNA sequences of 200 bp embedded with one SPI1 and one CEBPB motif as positive sequences. Negative sequences were then derived from this set by switching single nucleotides of the SPI1 motif for half of the positive sequences. Table 1 shows the top motifs output from MAGGIE and two comparative approaches, HOMER and a linear model adapted from the idea of MMARGE. Both MAGGIE and the linear model identified SPI1 and its similar motifs as the most significant hits. On the contrary, motif enrichment algorithms represented by HOMER identified both SPI1 and CEBPB as significant using the default random backgrounds for both positive (“pos vs. bg” column) and negative sequences (“neg vs. bg” column). HOMER lacked the sensitivity to capture the mutated SPI1 motif when the negative sequences were specified as the background in order to find the differential motif enrichment between the positive and negative set (“pos vs. neg” column).

**Table 1.** Top motifs output from different motif analysis tools tested on the simulated datasets.

| Rank | MAGGIE | Linear model | HOMER –<br>pos vs. neg | HOMER –<br>pos vs. bg | HOMER –<br>neg vs. bg |
| --- | --- | --- | --- | --- | --- |
| 1 | **SPI1\***<br>(13) | **SPI1\***<br>(8.6) | **SPI1**<br>(3) | CEBPG*<br>(198) | CEBPG*<br>(195) |
| 2 | SPIC*<br>(4.7) | ETV6<br>(3.6) | ETV6<br>(1) | CEBPD*<br>(194) | CEBPB*<br>(183) |
| 3 | ETV6*<br>(4.6) | SPIC<br>(3.6) | SPIC<br>(1) | CEBPB*<br>(192) | CEBPD*<br>(181) |
| 4 | IRF1<br>(2.2) | TFEB<br>(2.9) | STAT1::STAT2<br>(1) | CEBPE*<br>(191) | CEBPE*<br>(178) |
| 5 | ELF5<br>(2.0) | T<br>(2.4) | IRF1<br>(1) | **SPI1\***<br>(177) | **SPI1\***<br>(127) |
| 6 | THAP1<br>(2.0) | TFE3<br>(2.3) | SPIB<br>(0) | CEBPA*<br>(116) | CEBPA*<br>(105) |

-log<sub>10</sub> p-values are shown in parentheses. (\*) indicates motifs that passed FDR < 0.05 after the Benjamini–Hochberg controlling procedure. As the true positive, SPI1 motif is highlighted in bold.

In order to evaluate the sensitivity of MAGGIE, we tested its performance when different fractions of sequences were mutated at their SPI1 motifs (i.e., signal-to-noise ratio (SNR)). For each value of the SNR ranging from 10% to 80%, we repeated simulation of sequences 20 times and aggregated p-values for SPI1 and CEBPB from the three approaches. MAGGIE consistently outperforms the other methods in identifying the mutated motif (Fig. 2) and not the unmutated motif (Supplementary Fig. S2). Even though the other methods could potentially pass the significance threshold at a higher SNR or using a larger sample size (Supplementary Fig. S3), the true signal was not captured by these methods when motifs are mutated in less than 40% of the finite samples.

**Fig. 2.**
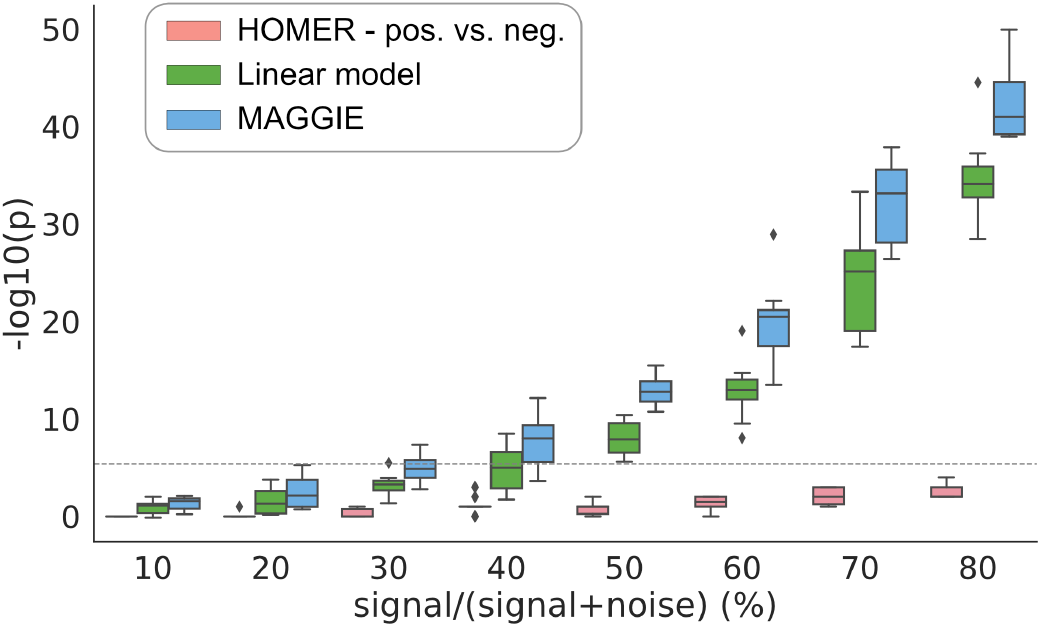
Comparison of sensitivity between MAGGIE and other approaches on simulated datasets. Significance values from three comparative approaches were shown for different pre-defined signal-to-noise ratios during data generation. Each boxplot aggregates the results from 20 simulations. During each simulation, a thousand sequences were randomly generated and inserted with SPI1 and CEBPB motifs, serving as the positive set. 10%-80% of these sequences experienced the change of a single nucleotide within their SPI1 motif, while the rest were kept untouched, resulting in the negative set. The same sets of positive and negative sequences were analyzed by the three approaches to facilitate the comparison. The dashed line indicates the significance threshold after multiple testing correction.

### 3.2 MAGGIE identifies known mediators for transcription factor binding sites and QTLs

After observing the superior sensitivity and specificity of MAGGIE on simulated data, we tested our method with several biological datasets. First, we analyzed the allele-specific TF binding sites caused by SNPs (Shi et al., 2016). Among the thirteen experiments tested, MAGGIE identified the corresponding motifs of the bound TFs for all of them (Fig. 3A). Even though p-value varies due to the quality and the sample size of each dataset, the corresponding motifs were recognized as the most significant even for TFs with as few as 37 allele-specific binding sites like USF1.

**Fig. 3.**
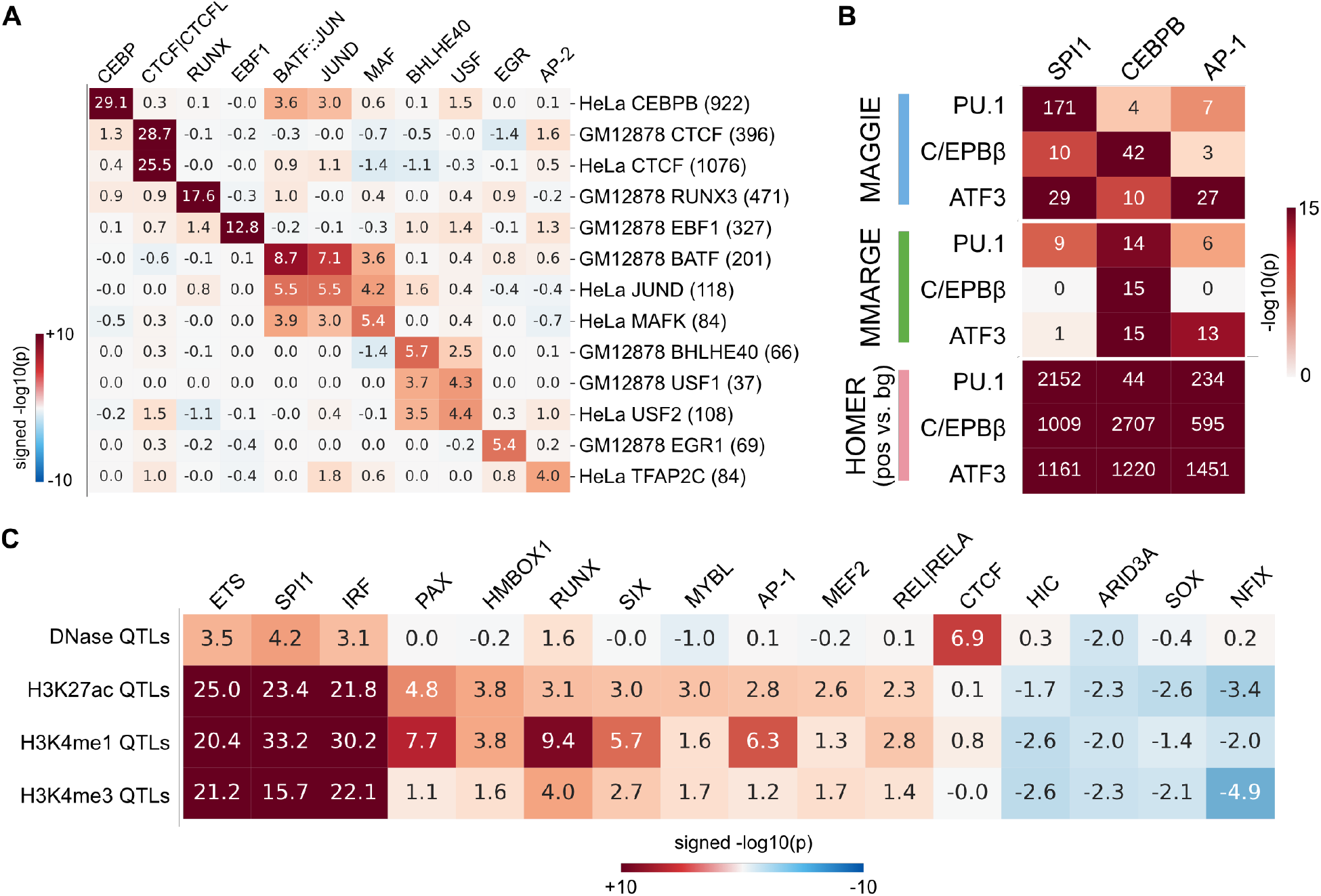
TF binding motifs identified by MAGGIE for various epigenomic features using biological datasets. (**A**) Signed significance values for allele-specific TF binding sites. Thirteen datasets were analyzed covering 12 different TFs from two cell types: GM12878 and HeLa-S3 (HeLa). Experiments are arranged vertically, and motifs are shown horizontally by their gene names. The sample size for each dataset is displayed in brackets. (**B**) Results for strain-specific TF binding sites. The log-transformed p-values for the three lineage-determining transcription factors (PU.1, C/EBPβ, ATF3) were shown and compared to the outputs from MMARGE and HOMER (positive sequences vs. random backgrounds). (**C**) Significant motifs identified for chromatin QTLs from lymphoblastoid cell lines. Motifs shown here were called significant for at least one type of the QTLs. The motifs are ordered by the signed significance values of H3K27ac QTLs. For all the results in this figure, results for similar motifs are averaged and displayed by their family names (e.g., CEBP, AP-1) or separated by “|”.

Next, we evaluated whether MAGGIE is able to recover the collaborative binding between TFs. Regulatory elements are usually bound by multiple TFs together, which form a complex with other co-activators to regulate functions (Reiter et al., 2017). For example, lineage-determining transcription factors (LDTFs) of macrophages such as PU.1, C/EBP, and AP-1 factors were frequently found to co-bind at macrophage-specific enhancers (Glass and Natoli, 2016; Heinz et al., 2015). Previous studies showed that the binding of specific LDTFs was not only dependent on each factor’s own motif, but also on nearby motifs recognized by collaborative factors (Heinz et al., 2013; Link et al., 2018a). To verify this conclusion with MAGGIE, we downloaded ChIP-seq data for PU.1, C/EBPβ, and ATF3, one of the AP-1 factors, from two genetically diverse strains of mice, C57BL/6J (C57) and BALB/cJ (BALB) (Link et al., 2018a; Fonseca et al., 2019). Strain-specific TF binding sites were identified for each factor and analyzed with MAGGIE. Sequences associated with TF binding were labeled as positive and their counterparts from the other strain as negative. We also used MMARGE to fit a linear mixed model between all the TF binding sites and their ChIP-seq read counts and used HOMER to find enriched motifs among positive sequences in comparison to random backgrounds. The outputs from the three approaches for the corresponding motifs are summarized in Figure 3B. MAGGIE recognized PU.1 binding to be mostly dependent on the PU.1 (or SPI1) motif instead of any other motif, while C/EBPβ binding was highly dependent on its own motif but also significantly dependent on SPI1 motif. The results were consistent with the pioneer role of PU.1 in opening chromatin and guiding the binding of other TFs (Barozzi et al., 2014). On the contrary, the comparative methods failed to distinguish the different functions between the bound TF and its collaborative factors. HOMER assigned strong significance to all the motifs because it was designed to identify enriched motifs without considering functions. MMARGE showed a lack of power in detecting collaborative factors using the two strains as it requires more data or larger genetic difference to confidently identify a linear relationship in the data.

The general framework of MAGGIE can also be applied to QTL datasets for epigenomic features that may be influenced by TF binding. We downloaded QTLs of several epigenomic features for lymphoblastoid cell lines (LCLs), including DNase I sensitivity quantitative trait loci (dsQTLs) for chromatin accessibility (Degner et al., 2012) and histone quantitative trait loci (hQTLs) for three types of histone modifications, H3K27ac, H3K4me1, and H3K4me3 (Grubert et al., 2015). Alleles are extended to 100-bp sequences and labeled as positive if associated with higher levels of the epigenomic features or negative if associated with lower levels. MAGGIE identified motifs with different specificity for the testing features (Fig. 3C). CTCF was output at top for dsQTLs but was insignificant for each type of hQTLs, supporting the major role of CTCF in maintaining chromatin structures instead of inducing active chromatin states (Arzate-Mejía et al., 2018). PU.1 together with other ETS factors were significant for both chromatin accessibility and histone modifications, indicating a pioneer role in opening chromatin as well as an important role in activating regulatory elements in LCLs (Scott et al., 1994). MAGGIE also identified many other motifs for histone modifications, which were found to maintain the cell identity and function of LCLs from previous studies, such as PAX factors (Glimcher and Singh, 1999), RUNX factors (Mevel et al., 2019), NF-κB factors (“REL|RELA”; Nagel et al., 2014). It is intriguing that several motifs showed up with potentially inhibitive functions, although these will need to be confirmed in later studies.

### 3.3 MAGGIE captures divergent functions of NF-κB factors for the stimulus responses of regulatory elements

Next, we tested MAGGIE with a more complex epigenomic feature: stimulus responses of regulatory elements. ATAC-seq and H3K27ac ChIP-seq data from four genetically diverse strains of mice were downloaded for macrophages at basal conditions and at pro-inflammatory conditions induced by 1-hour treatment of KLA (Link et al., 2018a). We used ATAC-seq data to locate open chromatin regions accessible for TF binding and H3K27ac ChIP-seq data to quantify the activity of these regions and identify active regulatory elements (Supplementary Fig. S4). By filtering for 2.5-fold change of activity from basal to KLA-treated conditions, we identified KLA-activated and KLA-repressed regulatory elements for each mouse strain (Fig. 4A). Among those, approximately 12000 activated elements and 18000 repressed elements were specific to one of the strains based on pairwise comparisons. Strain-specific activated and repressed regulatory elements were separately tested by MAGGIE to identify their mediators. Interestingly, besides PU.1 (or SPI1), CEBP, and AP-1 (or FOS::JUN) motifs that were known to be important for the KLA responses of macrophages (Glass and Natoli, 2016), MAGGIE assigned divergent functions for NF-κB factors (Fig. 4B). RELA corresponding to p65 subunit was output as functional for the activation response, while NFKB1 corresponding to p50 subunit was found significant for the repressed elements. On the contrary, the enrichment of these factors in activated or repressed elements was indistinguishable using HOMER due to the similarity of their motifs (Supplementary Fig. S5). Previous studies have shown that p65 frequently forms heterodimers with p50 to act as a transcriptional activator, while p50 homodimers result in transcriptional repression (Brignall et al., 2019; Cheng et al., 2011; Natoli et al., 2005). However, the genome-wide functions and binding patterns of these factors remain unknown.

**Fig. 4.**
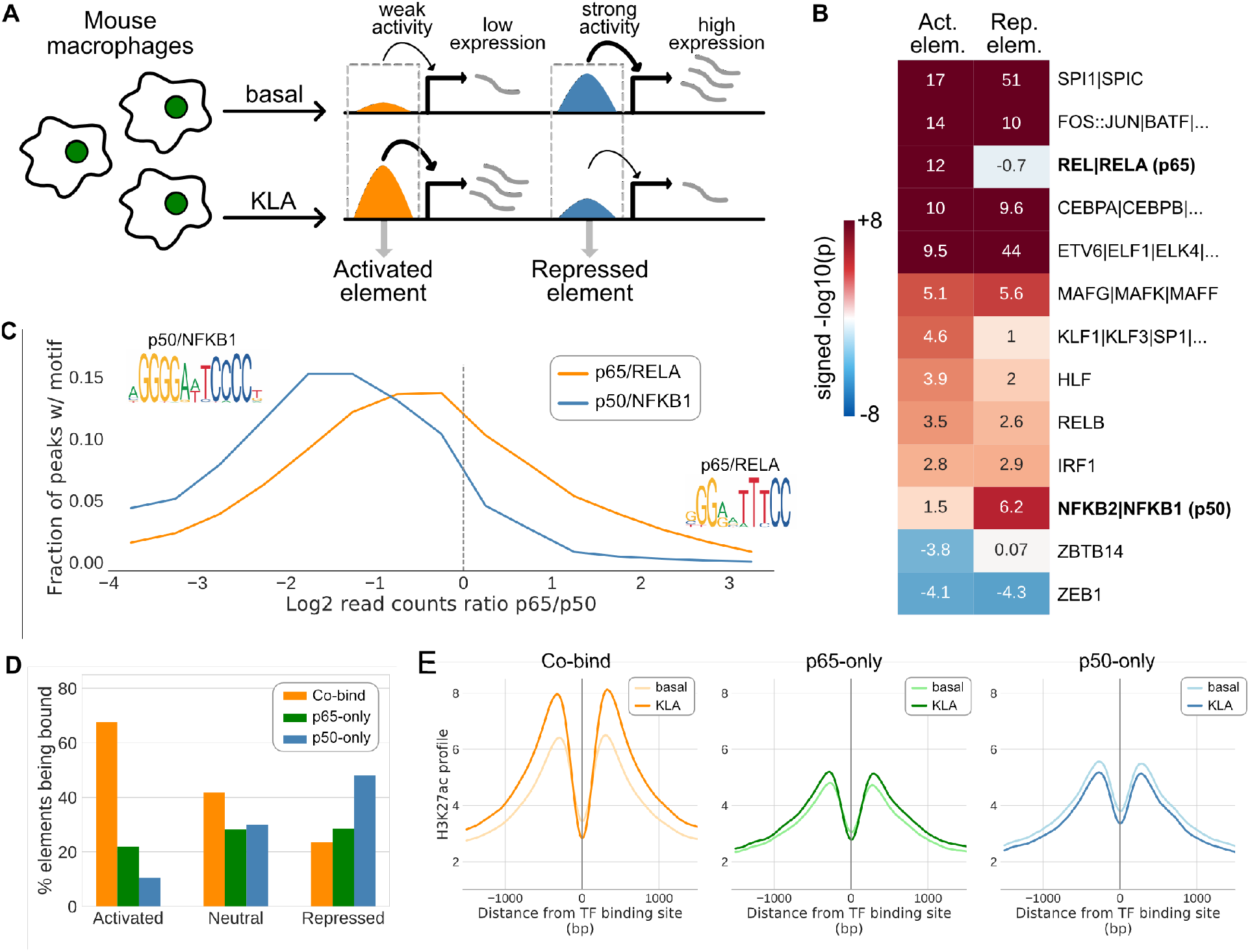
Divergent functions of distinct NF-κB factors in pro-inflammatory macrophages captured by MAGGIE and validated by experiments. (**A**) Sketch of KLA-activated and KLA-repressed regulatory elements defined by greater than 2.5-fold changes of H3K27ac from basal to KLA-treated conditions. (**B**) Significant results from MAGGIE on activated and repressed regulatory elements. Motifs are displayed by their gene names. Similar motifs are separated by “|” and shown with their average results. NF-κB factors are highlighted in bold. Protein names of RELA and NFKB1 are shown in the brackets, corresponding to p65 and p50, respectively. (**C**) Binding activities of NF-κB factors at sites with RELA or NFKB1 motifs. Motifs were searched within the 200-bp binding sites of p65 and p50 at the KLA-treated condition measured by ChIP-seq experiments. ChIP-seq reads for both p65 and p50 were counted to quantify binding activities. Regions with at least 32 reads of either factor were used to compute the log2 ratio of reads between p65 and p50. The distributions of log ratios are displayed in orange for sites having RELA motif (10549 sites) and in blue for sites having NFKB1 motif (2144 sites). The logos of motif PWMs are demonstrated as well. (**D**) Co-existence of NF-κB binding and the KLA responses of regulatory elements. NF-κB binding sites identified from ChIP-seq data were merged to find sites bound by p65 alone (p65-only), p50 alone (p50-only), or both (Co-bind). Among the regulatory elements that overlap with NF-κB binding sites, the bar plots summarized the fractions of elements bound by different NF-κB factors for activated, neutral, and repressed elements. (**E**) Change of H3K27ac at NF-κB binding sites after KLA treatment. H3K27ac ChIP-seq reads were counted within +/−1500 bp around the three categories of NF-κB binding sites using a bin size of 25 bp and were averaged to show the overall change of H3K27ac profiles at sites bound by different NF-κB factors.

To validate the functions of p65 and p50 for the KLA responses of macrophages, we conducted ChIP-seq experiments in C57 mice for p65 and p50 to measure their genome-wide binding sites in KLA-treated macrophages. Based on the measured TF binding sites, we first investigated the binding patterns of these factors. We searched for sites with RELA or NFKB1 motifs and computed the binding activities of NF-κB factors at those sites by counting ChIP-seq reads. Regions with relatively strong binding of either factor (greater than 32 ChIP-seq reads of p65 or p50; Supplement Fig. S6) were used to calculate the log ratios of read counts between p65 and p50 (Fig. 4C). RELA motif was enriched at the co-binding sites of p65 and p50, while NFKB1 motif was more strongly bound by p50. To connect the binding patterns to the regulatory elements used in MAGGIE, we overlapped the TF binding sites with the activated and repressed elements previously defined for C57 mice and found that the majority of activated elements were co-bound by both p65 and p50, while repressed elements were more often bound by p50 alone (Fig. 4D). By quantifying the regulatory activity around the binding sites of p65 and p50 by the level of H3K27ac, we found an overall decrease in H3K27ac around sites only bound by p50 and an overall increase around the co-binding sites of p65 and p50 after KLA treatment (Fig. 4E). These findings implicate a genome-wide role of p65-p50 heterodimers as a transcriptional activator and p50 homodimers as a repressor for KLA-treated macrophages. More importantly, our experimental results validated the outputs from MAGGIE regarding the divergent functions of p65 and p50 subunits for the KLA response of regulatory elements in macrophages, showing promise of using MAGGIE to discover novel functions of TFs for complex epigenomic features.

## 4. Discussion

We developed MAGGIE, a novel method for identifying DNA sequence motifs mediating TF binding and function. To our knowledge, MAGGIE is the first work to associate the mutation of TF binding motif with various types of epigenomic features. Our method focuses on the change of motif score and intentionally ignores the actual motif score due to the strong correlation between motif mutation and change of TF binding (Fig. 1C), and the independency of this relationship from the actual motif score (Supplementary Fig. S1). Another reason not to incorporate the actual motif score is that many epigenomic features do not possess a simple relationship with motif score. Linear models such as MMARGE and TBA worked well for TF binding but oversimplify the relationships for more complex features like histone modification or changes in the activity of regulatory elements. Instead, MAGGIE aggregates motif mutations associated with changes of epigenomic features and tests for a bias in the changing direction of mutations. We demonstrated that MAGGIE is able to identify known functional motifs for TF binding (Fig. 3A, B), chromatin accessibility (Fig. 3C), and histone modification (Fig. 3C). MAGGIE also helped to discover divergent functions of distinct NF-κB factors for the KLA response of regulatory elements in macrophages (Fig. 4), which was not found by any other motif analysis tools. It is worth noting that the motifs of NF-κB factors are usually too similar to be distinguished by motif enrichment algorithms (Supplementary Fig. S5), but the strategy of focusing on changes of motif score instead of the actual value of the motif score is sensitive enough to capture the difference.

MAGGIE was designed to take binary-labeled datasets as inputs (i.e., positive and negative sequences), facilitating application to most publicly available data. The framework can accept sequences derived from aggregated datasets like QTLs, or processed data from sequencing experiments like ChIP-seq and ATAC-seq. However, for the framework to work, MAGGIE requires additional measurements and genetic variation information for at least two different genotypes, which may not be currently available for some biological problems. Another potential limitation is the inevitable cutoff when assigning binary labels. We recommend analyzing with different cutoffs, especially when there are concerns about insufficient sample size or low data quality.

Theoretically, MAGGIE can be applied to any type of epigenomic feature that is potentially affected by TF binding. It will be interesting to investigate the performance of MAGGIE in other features, such as chromatin interaction and DNA methylation. Another future extension of MAGGIE is to assist with the identification of causal variants from GWAS. For instance, we can assign weights to the changes of motif scores caused by a variant based on the corresponding significance values coming out of MAGGIE and use these new metrics to prioritize variants. Given the high level of interest in understanding the function of genetic variation and the unprecedented generation of sequencing data for different individuals and animal strains, we expect MAGGIE, due to its flexibility for the type of input data, to be an effective bioinformatic tool that can be incorporated into the regular routine of motif analysis.

## Acknowledgements

We would like to express our great appreciation to Melissa Gymrek, Jenhan Tao, and Ludmil B. Alexandrov for assistance with manuscript editing. Our special thanks are extended to Inge R. Holtman for beta testing and feedback on the package, and Jana Collier for technical assistance.

