## Supplemental figures for "MAGGIE: leveraging genetic variation to identify DNA sequence motifs mediating transcription factor binding and function"

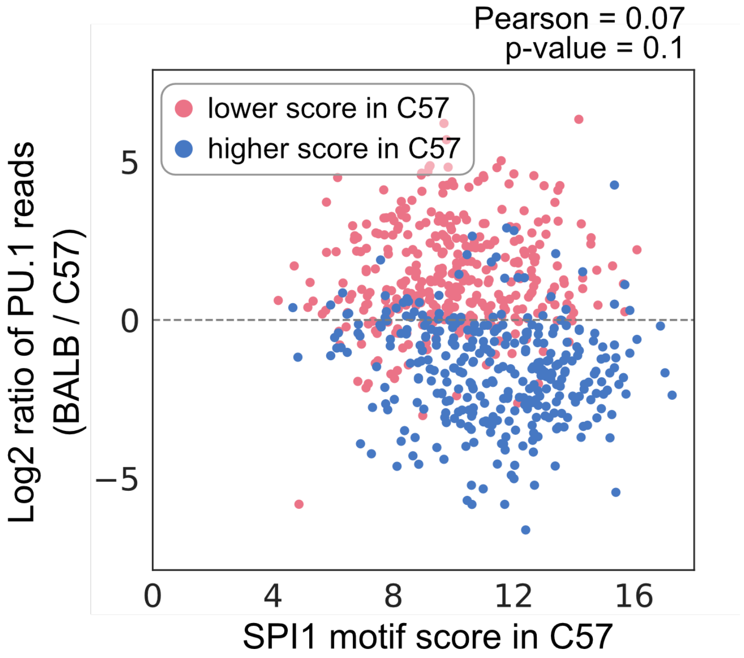


**Supplementary Fig. S1.** Relationship between change of PU.1 binding and SPI1 motif score at a similar level of SPI1 motif mutation. Each dot represents a PU.1 binding site that has mutation on SPI1 motif between BALB and C57 by a difference of motif score between 1 and 1.5. Red dots are binding sites with a lower motif score in C57, while blue dots are for sites with larger motif scores. Change of PU.1 binding activity was calculated by the fold change of PU.1 ChIP-seq reads between BALB mice and C57 mice. Sites with a stronger PU.1 motif in BALB has an increase in PU.1 binding in general, but the level of binding increase is not affected by the actual motif score (Pearson coefficient = 0.07 with an insignificant correlation).

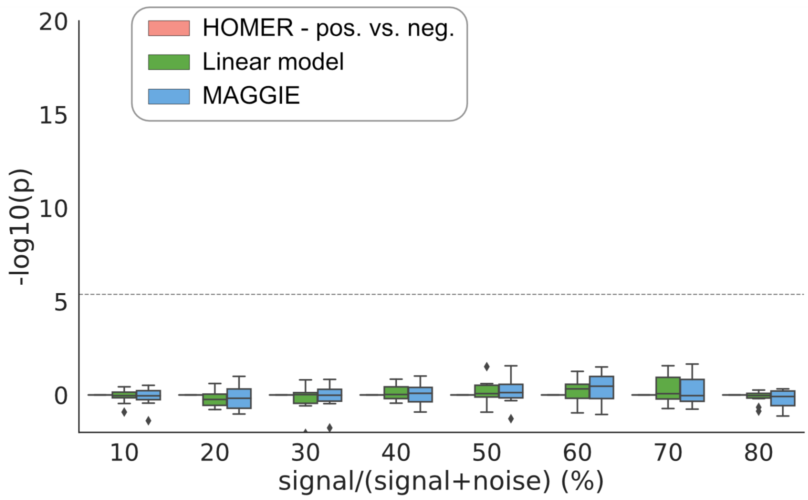


**Supplementary Fig. S2.** Significance values from the three comparative approaches for CEBPB motif on the simulated datasets. All of the methods recognize CEBPB motif as the un-mutated motif for all levels of simulated signal-to-noise ratio. The simulated datasets used here were the same as Fig. 2.

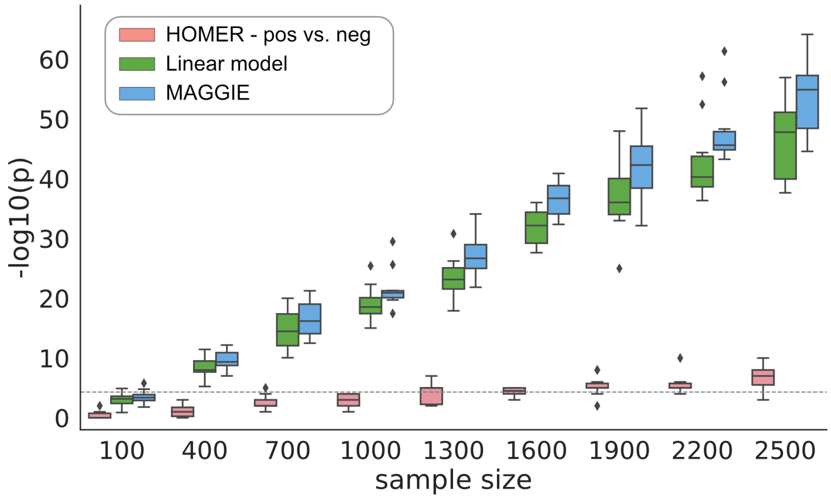


**Supplementary Fig. S3.** Effect of sample size on the outputs from the three comparative approaches. Different number of input sequences were simulated and embedded with SPI1 and CEBPB motifs, among which 50% of total sequences experienced mutation on the SPI1 motifs. Ten experiments were repeated for each sample size. Overall, significance values from all the three methods increased with a larger sample size.

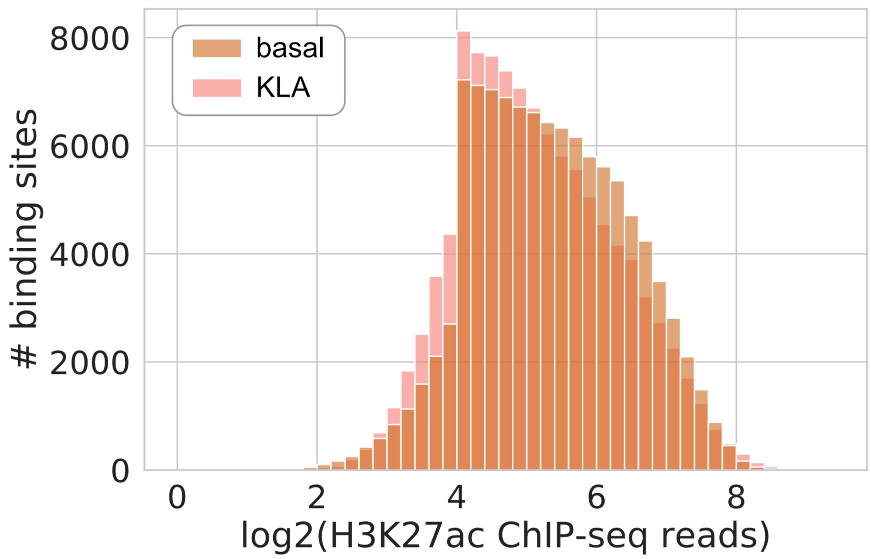


**Supplementary Fig. S4.** Distribution of H3K27ac ChIP-seq reads for extended open chromatin of macrophages at basal and KLA-treated conditions. ChIP-seq reads were counted within 1000-bp extended regions around open chromatin regions identified from ATAC-seq. Read counts were pooled for the four testing strains of mice. Discontinuity of read counts occurred at roughly 4 = log2(16), suggesting that regions with larger than 16 reads are more confidently called as active regulatory elements.

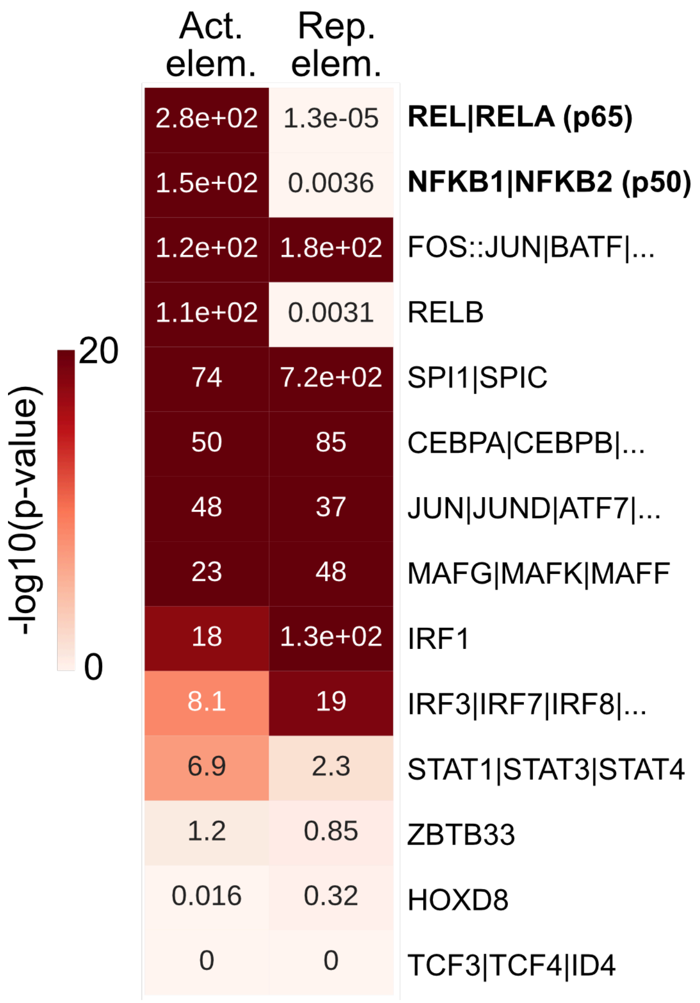


**Supplementary Fig. S5.** HOMER results for KLA-activated and KLA-repressed regulatory elements. Motif enrichment for known motifs was calculated using HOMER with default parameters. RELA motif was found overrepresented in both activated and repressed regulatory elements, while NFKB1 motif did not show significant enrichment in the whole set of those elements but was only identified as a functional motif for repressed elements by leveraging genetic variation data.

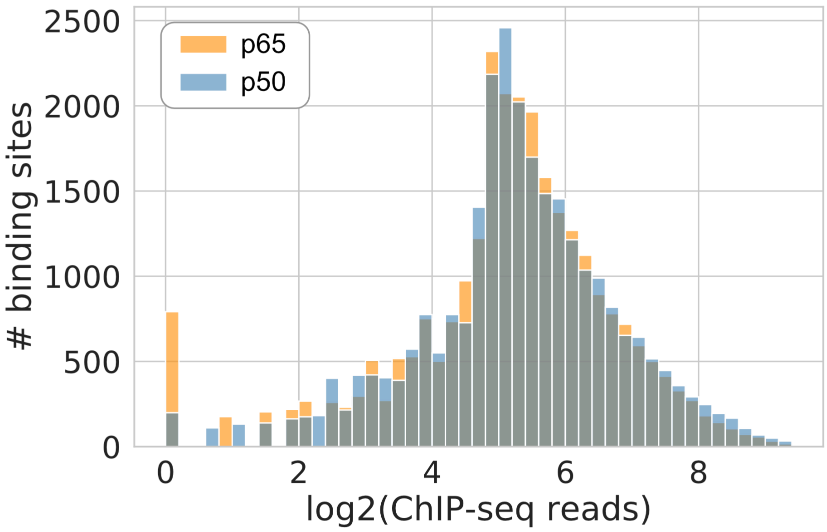


**Supplementary Fig. S6.** Distribution of ChIP-seq reads for p65 and p50 at their respective binding sites. ChIP-seq reads were counted within 200-bp binding sites called for p65 and p50 using HOMER “findpeaks -size 200”. The median counts for both factors are roughly at 5 = log2(32 reads).
